# An ATP-binding cassette subfamily C is crucial for flavonoid sequestration in the domestic silkworm, *Bombyx mori*

**DOI:** 10.1101/2025.05.14.652641

**Authors:** Ryusei Waizumi, Chikara Hirayama, Kenji Watanabe, Tetsuya Iizuka, Yoko Takasu, Hideki Sezutsu

## Abstract

Some herbivorous insects have evolved sequestration mechanisms, the ability to take up and accumulate plant secondary metabolites for their own benefit. The domestic silkworm, *Bombyx mori*, and its wild ancestor, *B. mandarina*, take up flavonoids from mulberry leaves and accumulate the molecules as their glucosides in their tissues and cocoon shell. This sequestration enhances the cocoon’s protective property against ultraviolet or bacterial proliferation. Here, we show that an ATP-binding cassette transporter subfamily C (ABCC) gene, *BmABCC4*, plays a crucial role in the flavonoid sequestration in the silkworms. *BmABCC4* is located at *Green c*, a cocoon color-associated locus predicted in 1941 and whose detailed position we previously identified. This transporter is expressed in the midgut and silk glands, and the expression is upregulated in the midgut in late period of final instar larva. Knockout of *BmABCC4* significantly reduced the total flavonoid content in the tissues and cocoon shell. Our results suggest that BmABCC4 transports flavonoid glucosides from the midgut cells to hemolymph and from the silk gland cells to the silk gland lumen.

## Introduction

Plants have evolved to synthesize various chemical compounds to deter herbivory through co-evolution with insects. However, some insects have not only evolved mechanisms to detoxify these defense compounds but also to accumulate them for their own defense. This strategy, known as sequestration, plays a key role in the evolution of plant-herbivore interactions. Flavonoid is a representative class of plant secondary metabolites. Although not highly toxic, flavonoids can strongly deter insect feeding and oviposition, and inhibit growth, depending on their concentration (Simmonds 2003). However, some lepidopteran and orthopteran insects are known to extensively sequester flavonoids in their tissues (Hirayama *et al*., 2013; Hopkins and Ahmad 1991; Schittko *et al*., 1999). The domestic silkworm, *Bombyx mori*, and its wild ancestor, *Bombyx mandarina*, are instances of such insects; they sequester quercetin and kaempferol from mulberry leaves into their tissues and cocoon shell (Fujimoto and Hayashiya 1961; Hirayama *et al*., 2009). The flavonoids accumulated in the cocoon shell exhibit antimicrobial properties and play a crucial role in protecting tissues from ultraviolet radiation during pupation (Daimon *et al*., 2010; Hirayama *et al*., 2008; Kurioka *et al*., 1999). Understanding the molecular mechanisms of flavonoid sequestration in the silkworms could provide significant insights into plant-herbivore interactions and co-evolution.

Since sequestered flavonoids color the cocoon shell yellow-green, a trait that attracts agronomic interest, many attempts have been made over the past century to identify cocoon color-associated loci through classical linkage analysis. These loci have conventionally been named with terms including “Green.” In other words, these efforts aimed to identify genes involved in flavonoid sequestration. Recent studies have identified genes involved in flavonoid metabolism, elucidating several steps of flavonoid sequestration from uptake into the midgut to accumulation in the silk: (i) quercetin uptake into midgut cells mainly depends on the deglycosylation of its glycosides by the hydrolytic enzyme Glycoside hydrolase family 1 group G 5 (GH1G5), encoded at the locus *Green d* (Waizumi *et al*., 2024); (ii) the imported quercetin is re-glucosylated at the 5-*O* position by uridine 5’-diphospho-glucosyltransferase (UGT), encoded at the locus *Green b* (Daimon *et al*., 2010); and (iii) the flavonoid glucosides are likely transported from the hemolymph into silk gland cells by the sugar transporter StrGn2-7, encoded at the locus *New Green Cocoon* (Lu *et al*., 2023). In our previous study, we identified a quantitative trait locus (QTL) associated with flavonoid content in the cocoon shell on chromosome 15 (Waizumi *et al*., 2024). In 1941, Hashimoto predicted a locus named *Green c* (*Gc*), associated with the yellow-green color of cocoons (Hashimoto 1941). Subsequently, it was predicted to be located at an unknown position on the 15th linkage group (Fujii *et al*., 1998; Fujimoto 1955). Considering these studies, we deemed the QTL identical to *Gc*. However, no genes involved in the flavonoid sequestration have been identified around this position. Identifying the unknown associated gene is expected to provide significant insights into the understanding of insect sequestration strategies.

Here, we report that an ATP-binding cassette transporter subfamily C member 4 (BmABCC4) plays a crucial role in flavonoid sequestration in the silkworms. Since *BmABCC4* is located at the QTL on chromosome 15, the prediction of *Gc* as a cocoon color-associated locus has been confirmed. Our results suggest that BmABCC4 mediates flavonoid glucoside transport from midgut cells to hemolymph, and from silk gland cells to the silk gland lumen. This discovery fills a crucial gap in understanding flavonoid sequestration in the silkworms.

## Materials and methods

### Transmembrane domain prediction

We integrated prediction results from TMHMM v2.0 and SOSUI v1.11 because these tools predicted different positions of domains in ABCC4 (TMHMM and SOSUI respectively predicted 1st–6th domains and 7th-12th) (Hirokawa et al., 1998; Krogh *et al*., 2001). The nucleotide binding domains (NBDs) were predicted through multiple alignment with known NBD sequences (Oswald *et al*., 2006). The sequence feature of ABCC4 was visualized using Protter v1.0 (https://wlab.ethz.ch/protter/start/).

### Insect materials

The silkworm strain p50T is maintained at the National Agriculture and Food Research Organization (NARO, Japan). The silkworm eggs were incubated under 15-h light/9-h dark photoperiod until hatching, and the larva were reared on a commercial artificial diet (SilkMate PM; Nosan, Kanagawa, Japan) under 12-h light/dark photoperiod at 25°C. All individuals were female except those from which RNA-seq data were derived.

### Expression analysis

Total RNA was extracted using ISOGEN according to the manufacturer’s instruction. After DNA digestion and purification by DNeasy (Qiagen, Hilden, Germany), cDNA was synthesized using ReverTra Ace ® qPCR RT Master Mix (TOYOBO, Osaka, Japan). Quantitative RT-PCR reactions were performed with Luna Universal qPCR Master Mix (New England Biolabs, Ipswich, Massachusetts, U.S.A.) on LightCycler® 96 system (Roche, Basel, Switzerland). Data were normalized to *rp49* (SilkBase gene ID: *KWMTBOMO14639*) by the comparative CT (2-DDCT) method. The sequences of primers used are listed in S1 Table. RNA-seq data from third-day fifth instar larvae were previously published by our research group (Waizumi *et al*., 2023; Yokoi *et al*., 2021). The sequencing reads were trimmed by fastp v0.20.0 (Chen *et al*., 2018) with the following parameters: -q 20 -n 5 -l 100. Salmon v1.5.2 (Patro *et al*., 2017) was used to calculate read counts and transcripts per million (TPM) scores. The published gene model for p50T was used as the mapping reference (Kawamoto *et al*., 2019).

### Flavonoid content measurement

The flavonoid content in cocoons was measured using a simplified method that utilizes the high correlation between flavonoid content and the absorbance at 365 nm in methanol extracts from cocoon fragments (Hirayama and Okada 2014). The extraction was conducted using shredded 2– 3-mm squares of the cocoons with MeOH–H2O (7:3, V/V) at 60°C for 2 h and repeated for two times for thorough extraction. The flavonoid content in tissues was measured using a HPLC system (LC-10A, Shimadzu Co., Kyoto, Japan) according to a previous report (Waizumi *et al*., 2024). In brief, contents were separated SunFire C18 column (150 × 3.0 mm i.d.; Waters, Massachusetts, USA) at a flow rate of 1.0 mL/min and detected by n SPD-M10AVP photodiode array detector (Shimadzu Co.). Flavonoids were identified based on their distinctive biphasic spectral characteristics, peaking at around 260 and 380 nm. Quercetin 5, 4 ’-di-*O*-glucoside was used as the reference compound.

### Genome editing

The TALEN-mediated genome editing was conducted according to a previous report (Takasu *et al*., 2013), using a Golden Gate TAL and TALEN Kit (Cermak *et al*., 2011). TALEN mRNAs were microinjected into eggs of the p50T strain within 8 h after oviposition. The eggs were treated with a corona discharge just after oviposition to prevent embryonic diapause (Yamada *et al*., 2022). The mutant lines were established by high-resolution melting analysis-mediated genotyping and mating as described previously (Waizumi *et al*., 2024). Sanger sequencing determined the mutated sequences. The sequences of primers used are listed in S1 Table.

### Phylogenetic analysis

The homologous proteins were collected by BLASTp search using *Gc*-coded 5 ABCC proteins as the query. Proteins with e-values less than 1E-100 were used for the analysis. Protein models of 10 lepidopteran insects available in InsectBase 2.0 (Mei et *al*., 2022) were used for the BLAST database. *Arabidopsis* ABC subfamily B member 1 (AtABCB1) was used as an outgroup. Proteins were aligned using Clustal Omega v1.2.4 (Sievers *et al*., 2011). The phylogenetic tree of lepidopteran orthologs of the *Gc*-coded ABCC proteins was constructed by the maximum likelihood method using RAxML-NG v1.1 (Kozlov *et al*., 2019) with the following parameters: -bs-trees 100, -model LG+I+G4+F. The tree was visualized using iTOL v5 (Letunic and Bork 2021). A species tree of Lepidoptera was drawn regarding the previous report (Kawahara *et al*., 2019) (S1B Fig).

## Results

### *BmABCC4* was identified as the *Gc*-coded candidate gene involved in the flavonoid sequestration

In our previous study, we conducted a QTL analysis focused on flavonoid content in the cocoon shell using F_2_ progenies of two different domestic silkworm strains, p50T (forming flavonoid-abundant yellow-green cocoon) and J01 (forming flavonoid-deficient white cocoon) (Waizumi *et al*., 2024). Among the identified three QTLs, one on chromosome 15 was considered to be *Green c*, a cocoon color-associated locus previously predicted on the 15th linkage group without detailed position information (Fujii *et al*., 1998; Fujimoto 1955; Hashimoto 1941). The flavonoid content in the cocoon shells of the J01 type of *Gc* (+/+) was about 70% of that in the p50T strain type (*Gc*/*Gc*), when the genotypes of the other two QTLs (*Gd* and *Gn*) were heterozygous of p50T and J01-types (Fig 1A). Near the closest marker to the QTL corresponding to *Gc* (Chr15: 9446209 bp), a cluster of five genes encoding ABC transporters was located (KWMTBOMO08968: 8977053–9010926 bp, KWMTBOMO08969: 9015657–9033392 bp, KWMTBOMO08974: 9088293–9114754 bp, KWMTBOMO08988: 9230273–9271571 bp, KWMTBOMO09024: 9884645–9916355 bp) (gene IDs are from SilkBase: Kawamoto *et al*., 2019). Given that this transporter family is indicated to be involved in the transport of flavonoid glycosides in plants (Beherens *et al*., 2019; Francisco *et al*., 2013), we hypothesized that these genes are involved in flavonoid sequestration in the silkworms. All these clustered genes are classified as ABC subfamily C (ABCC) (Kawamoto *et al*., 2019). Among these, we focused on *KWMTBOMO08988* because its position is closest to the marker. A previous study has named this gene *BmABCC4* (Adegawa *et al*., 2024). BmABCC4 consists of 1363 amino acid residues and includes six transmembrane domains and a pair of NBDs, typical domains of ABC transporters. BmABCC4 in the J01 strain contained four amino acid substitutions (A723G, V785T, A1089V, P1092S) compared to the orthologous protein in the p50T strain (Fig 1B). We hypothesized that the substitutions include the causative mutation responsible for the differences in the cocoon shell flavonoid content between the strains.

**Fig 1.**
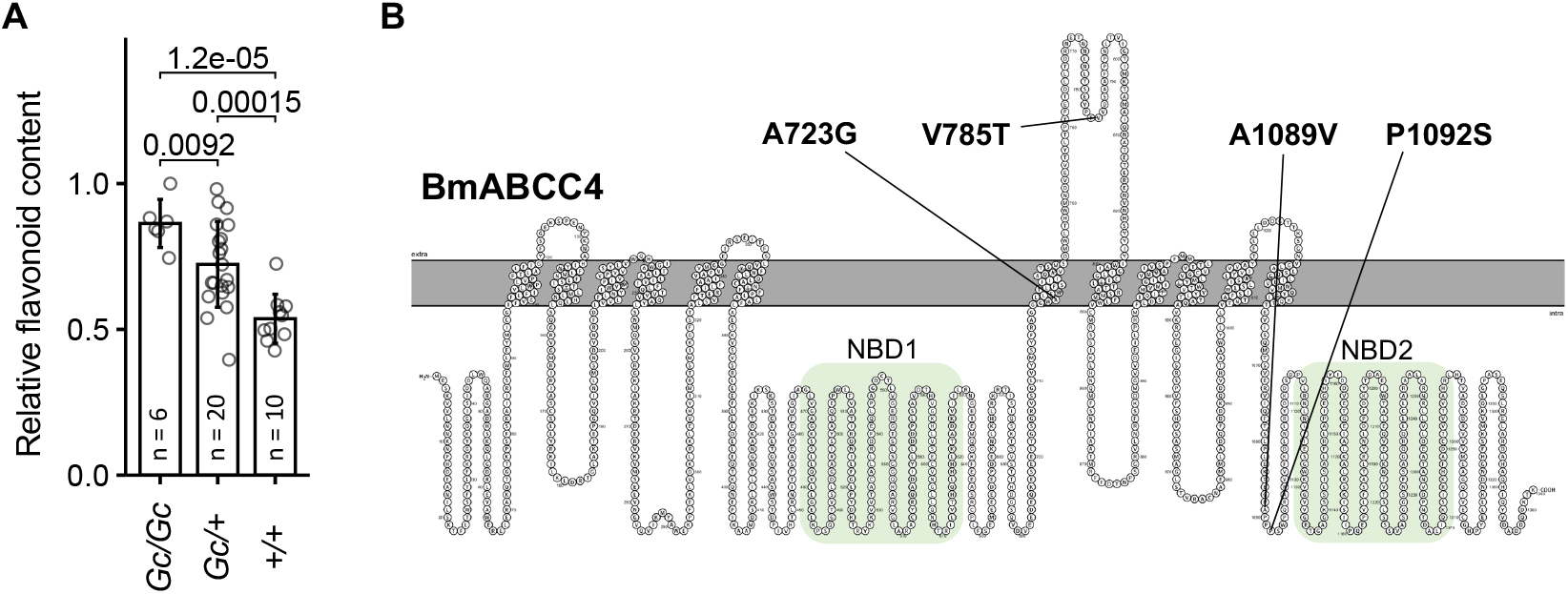
Identification of BmABCC4 as the *Gc*-coded candidate protein involved in the flavonoid sequestration. (A) Effect of *Gc* genotype on flavonoid content in the cocoon shell. Data are from phenotyping result of our previous report conducting QTL analysis focused on flavonoid content in the cocoon shell (Waizumi *et al*., 2024). All individuals from which the phenotype data are derived possess *Gd*/+ and *Gn*/+ genotypes. Data are means of n independent biological replicates ±SD. The values above the graph are p-values calculated by a two-tailed Student’s t-test. (B) Sequence feature of BmABCC4. Amino acid substitutions in J01 compared to p50T are indicated.

### *BmABCC4* is expressed in the midgut and silk glands in the final instar larva

One of the well-known function of ABC transporters in mammals is the extracellular export of molecules (Khunweeraphong and Kuchler 2021). Therefore, we hypothesized that BmABCC4 is involved in the export of flavonoids from the midgut cells to the hemolymph or from the silk gland cells to the lumen of the silk gland within the pathway from flavonoid uptake to accumulation in silk. Using published RNA-seq data of third-day fifth-instar larvae of p50T and J01 (Waizumi *et al*., 2023; Yokoi *et al*., 2021), we quantified the expression levels in the midgut and silk glands (Fig 2A). *BmABCC4* was strongly expressed in the midgut. Although at lower levels, it was also expressed in the silk glands, particularly in the anterior part of the middle silk glands. The lower expression levels in p50T compared to J01 in both the midgut and silk glands suggested that the differences in the cocoon shell flavonoid content between the strains are due to differences in the encoded sequences rather than the expression levels. Furthermore, we examined the expression levels of *BmABCC4* during fifth instar larval stage by qPCR (Fig 2B). Although no clear pattern was observed in the expression changes in the middle and posterior silk glands, the expression in the midgut was increased in the later period. This result was consistent with the rapid progression of flavonoid accumulation in the silk gland during the late fifth instar larval stage (Lu *et al*., 2023).

**Fig 2.**
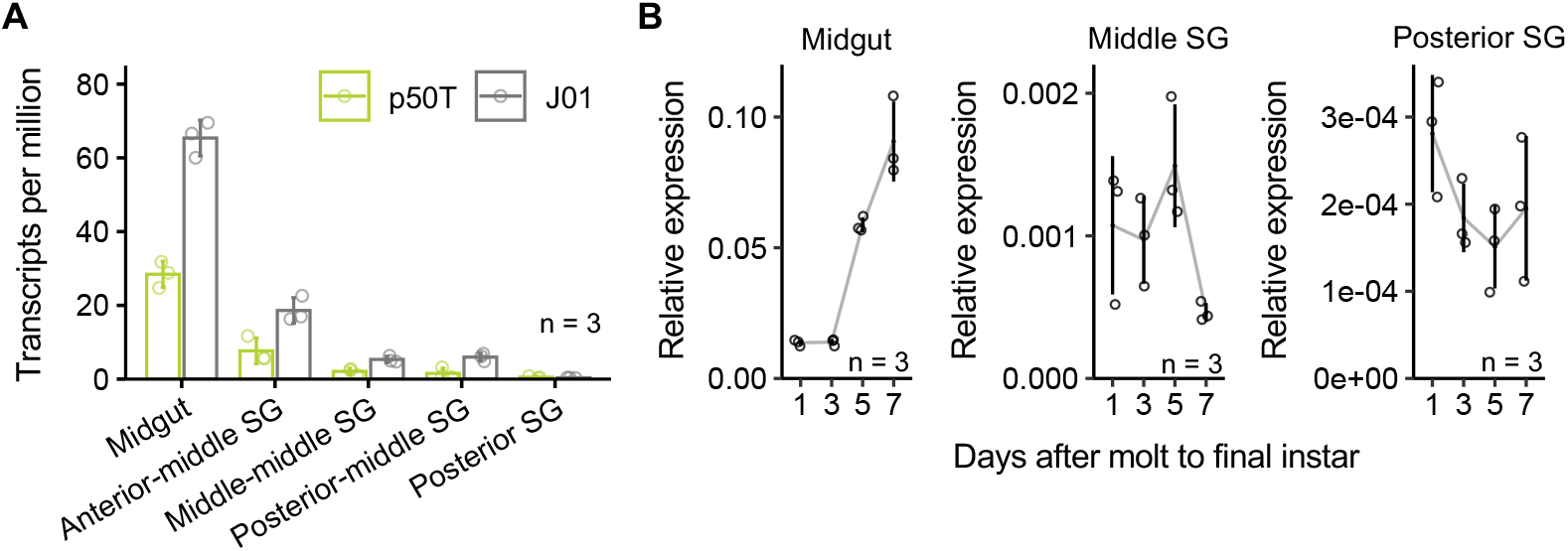
Expression pattern of *BmABCC4* in the larval organs responsible for the flavonoid sequestration. (A) Expression pattern of *BmABCC4* extracted from RNA-seq data of 3rd-day final instar larva of p50T and J01. (B) Expression changes of *BmABCC4* during fifth instar larval stage. Data are means of n independent biological replicates ±SD.

### Knockout of *BmABCC4* reduced flavonoid content in the organs and cocoon shell

To examine that BmABCC4 is involved in the flavonoid sequestration in the domestic silkworm, we produced knockout lineages of *BmABCC4* using transcription activator-like effector nucleases (TALEN). We obtained two knockout lineages of *BmABCC4* with different frameshift mutations in exon 4. We designated the one with a 4-bp deletion as *abcc4*^*Δ4*^ and the other with a 11-bp deletion as *abcc4*^*Δ11*^ (Fig 3A). The mutations resulted in a premature stop codon at exon 4 along with shortened amino acid sequence lengths from 1363 to 137 in *abcc4*^*Δ4*^, and to 123 in *abcc4*^*Δ11*^. The mutants produced clearly discolored cocoons compared to p50T (Fig 3B). In addition, we observed a reduction of fluorescence under ultraviolet irradiation in the hemolymph (Fig 3C); such fluorescence is characteristic of quercetin glucosides formed by glycosylation at the 5-*O*-position (Daimon *et al*., 2010). The flavonoid content in the cocoon shell, midgut, hemolymph, and silk glands was reduced to less than half due to the knockout of *BmABCC4* (Fig 3D). The reduction rate was smallest in the midgut, followed by the hemolymph, posterior silk gland, middle silk gland, and largest in the cocoon shell. These results indicated that the knockout of *BmABCC4* caused dysfunction in the transport of flavonoid glucosides, such as quercetin glucosides glycosylated at the 5-*O* position.

**Fig 3.**
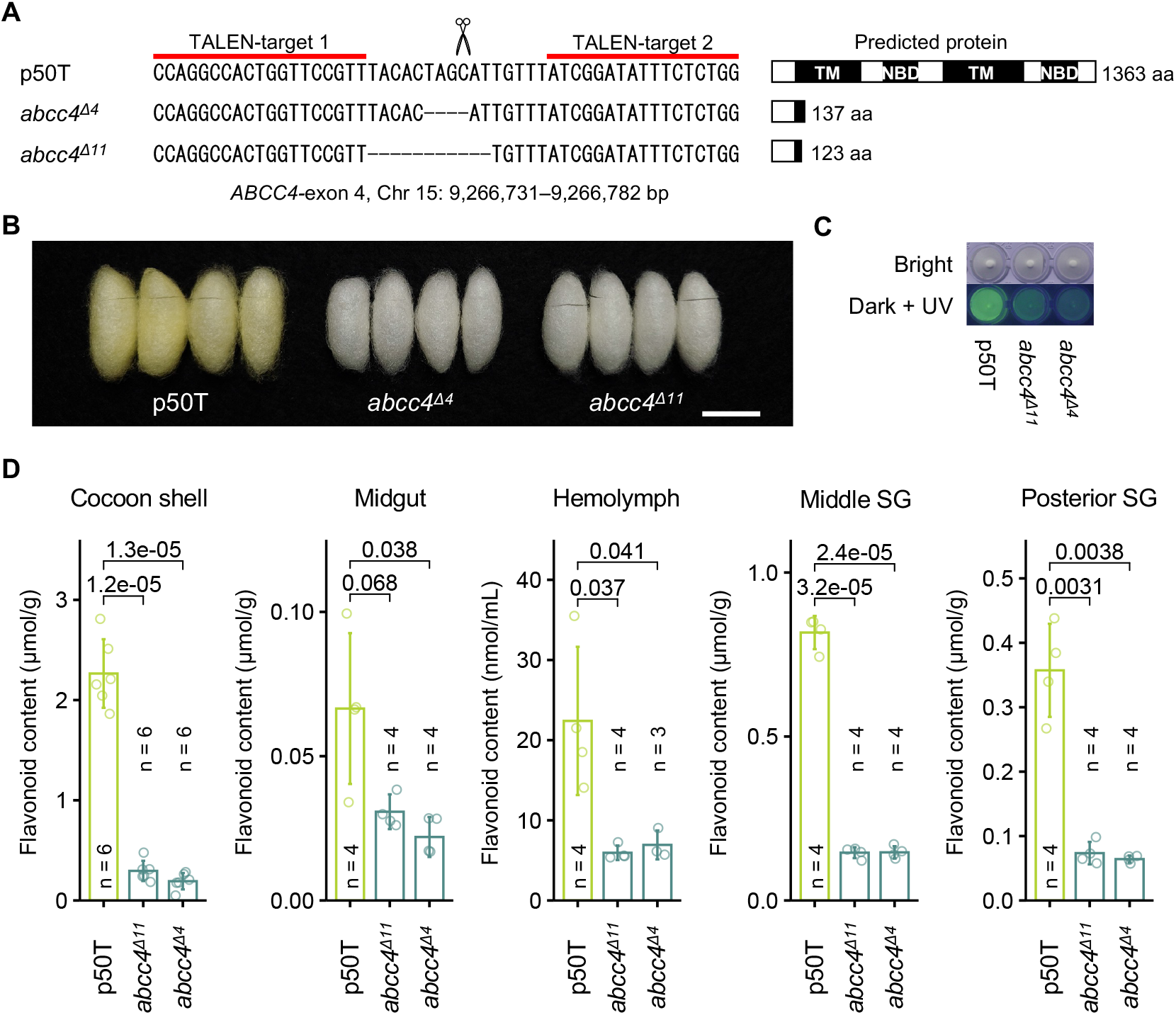
Reduction in flavonoid content in *BmABCC4*-knockout mutants. (A) Disrupted target sequences in *BmABCC4*. (B) and (C) Cocoons (B) and hemolymph (C) of the p50T strain and the *BmABCC4*-knockout mutant lineages. Bars = 20 mm. (D) Total flavonoid content of cocoon shells and tissues of the p50T strain and the *BmABCC4*-knockout mutant lineages. The values above the graph are p-values calculated by a two-tailed Student’s t-test. Data are means of n independent biological replicates ±SD. All insects were reared on a commercial artificial diet. SG, silk gland.

## Discussion

In this study, we showed that an ABCC plays a crucial role in flavonoid sequestration in the domestic silkworm. This is the first report indicating the involvement of ABC transporters in flavonoid sequestration in insects. *BmABCC4* is expressed in the midgut and silk glands of final instar larvae (Fig 2A). The increase in expression in the midgut during the late stage of the final instar larvae may be associated with the increase in flavonoid content in the silk glands during the same period (Lu *et al*., 2023) (Fig 2B). The expression in the silk glands is weaker compared to the midgut (Fig 2A and 2B). However, the cumulative reduction in flavonoid content in the responsible organs due to *BmABCC4* knockout suggests that *BmABCC4* is involved in flavonoid transport not only in the midgut but also in the silk glands; the reduction rate was lower in the midgut compared to the hemolymph, in the hemolymph compared to the silk glands, and in the silk glands compared to the cocoon shell (Fig 3D). By contrast, when *GH1G5*, which mediates flavonoid uptake in the midgut, is knocked out, the degree of restriction in flavonoid content in the midgut is carried over to all subsequent organs on the pathway of flavonoid transport (Waizumi *et al*., 2024). Our findings may have filled in the missing pieces of the major pathway of flavonoid sequestration in the domestic silkworm, from uptake in the midgut to accumulation in the silk. However, our results have raised a new question regarding the flavonoid transport mechanism. The knockout of *BmABCC4* resulted in a reduction of flavonoid content in the midgut to less than half (Fig 3D). This was contrary to our expectations; if BmABCC4 were localized on the basolateral side of the midgut cells and responsible for exporting flavonoids into the hemolymph, the knockout of *BmABCC4* would be expected to increase flavonoid content in the midgut due to the disruption of the export pathway. Vesicle-mediated transport may explain this result. In case of a sequestering leaf beetle, *Chrysomela populi*, an ABC transporter has a crucial role in sequestration of the phenolglucoside salicin. Since the transporter localizes at intracellular membrane of secretory cells, it is suggested that the transporter secretes salicin via a vesicle-mediated transport system rather than directly exporting it extracellularly (Strauss *et al*., 2013). If flavonoid transport in *Bombyx mori* midgut cells is also mediated by a vesicular transport involving BmABCC4, then the knockout of *BmABCC4* could disrupt the temporary storage of flavonoids in vesicles, potentially leading to a decrease in the cellular flavonoid content. To better understand the mechanism of flavonoid transport mediated by BmABCC4 in the domestic silkworm, it will be necessary to investigate the localization of BmABCC4 and the effect of *BmABCC4* knockout on the amount of flavonoid within vesicles.

Flavonoid sequestration is observed in lepidopteran insects other than *Bombyx* (Hirayama *et al*., 2013; Schittko *et al*., 1999). Given the commonness of flavonoid synthesis in vascular plants (Yonekura-Sakakibara *et al*., 2019), flavonoid sequestration may potentially occur even in numerous lepidopteran species where it has not been reported. In a maximum likelihood-inferred phylogenetic tree of lepidopteran-wide orthologous proteins of *Gc*-coded ABCC transporters, the ABCC4 clade includes homologs from all investigated species, including the most ancient lepidopteran family Micropterigoidea (S1A and S1B Fig). The subclades of each *Gc*-coded ABCC protein included most of the examined species belonging to Ditrysia (a group that emerged after Tineoidea) (S1A and S1B Fig). These results suggest that *Gc*-coded ABCCs have been functionally differentiated, with each protein playing an adaptively important role, and that ABCC4-mediated flavonoid sequestration is widely conserved in Lepidoptera. Therefore, the orthologs of BmABCC4 will be crucial clues for understanding the evolution of flavonoid sequestration in many lepidopteran insects and the interactions between lepidopteran insects and plants.

## Supplementary information

Supplementary materials are available at Figshare. doi: 10.6084/m9.figshare.28877480

## Data availability

Not applicable.

## Acknowledgements

We thank Ms. Mayuko Sakato for her technical assistance, and Sae Furuhashi, Toshihiko Misawa, Kaoru Nakamura, Keita Hiruma, Koji Hashimoto, and Eiji Okada for their support in the silkworm rearing. The computation was partly performed using the supercomputer of the Agriculture, Forestry and Fisheries Research Information Technology Center of the Japanese Ministry of Agriculture, Forestry and Fisheries.

## Author contributions

Conceptualization: Ryusei Waizumi, Chikara Hirayama, Tetsuya Iizuka.

Investigation: Ryusei Waizumi, Chikara Hirayama, Tetsuya Iizuka, Kenji Watanabe, Yoko Takasu

Supervision: Hideki Sezutsu.

Visualization: Ryusei Waizumi.

Writing – original draft: Ryusei Waizumi.

Writing – review & editing: Ryusei Waizumi, Chikara Hirayama, Tetsuya Iizuka, Kenji Watanabe,

Yoko Takasu

## Description of supplementary materials

**S1 Fig.**
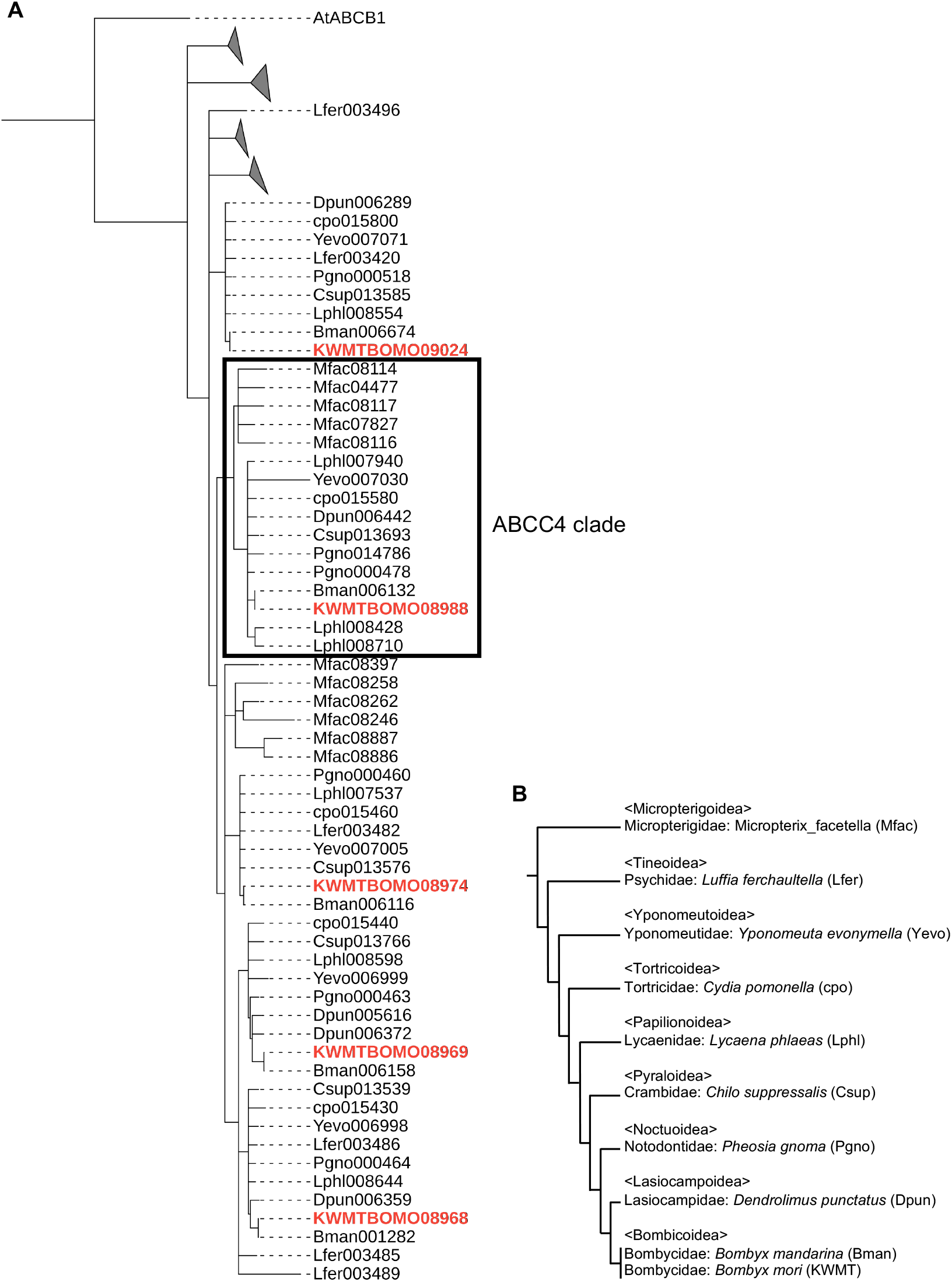
Phylogenetic tree of lepidopteran-wide orthologous proteins of *Gc*-coded ABCCs. (A) A maximum likelihood-inferred phylogenetic tree of lepidopteran-wide orthologous proteins of *Gc*-coded ABCC transporters. Unreliable nodes with bootstrap values (bootstrap trials = 100) under 90 are shown as multi-branching nodes. The tree was rooted using *Arabidopsis* ABCB1 as an outgroup. Labels of *B*.*mori Gc*-coded ABCCs are highlighted by red coloring. (B) Species tree of Lepidoptera drawn regarding a previous report (Kawahara *et al*., 2019)

S1 Table. Primers used in the experiments.

