## Supplementary figures and images for "An ATP-binding cassette subfamily C is crucial for flavonoid sequestration in the domestic silkworm, *Bombyx mori*"

### Supplementary figure S1

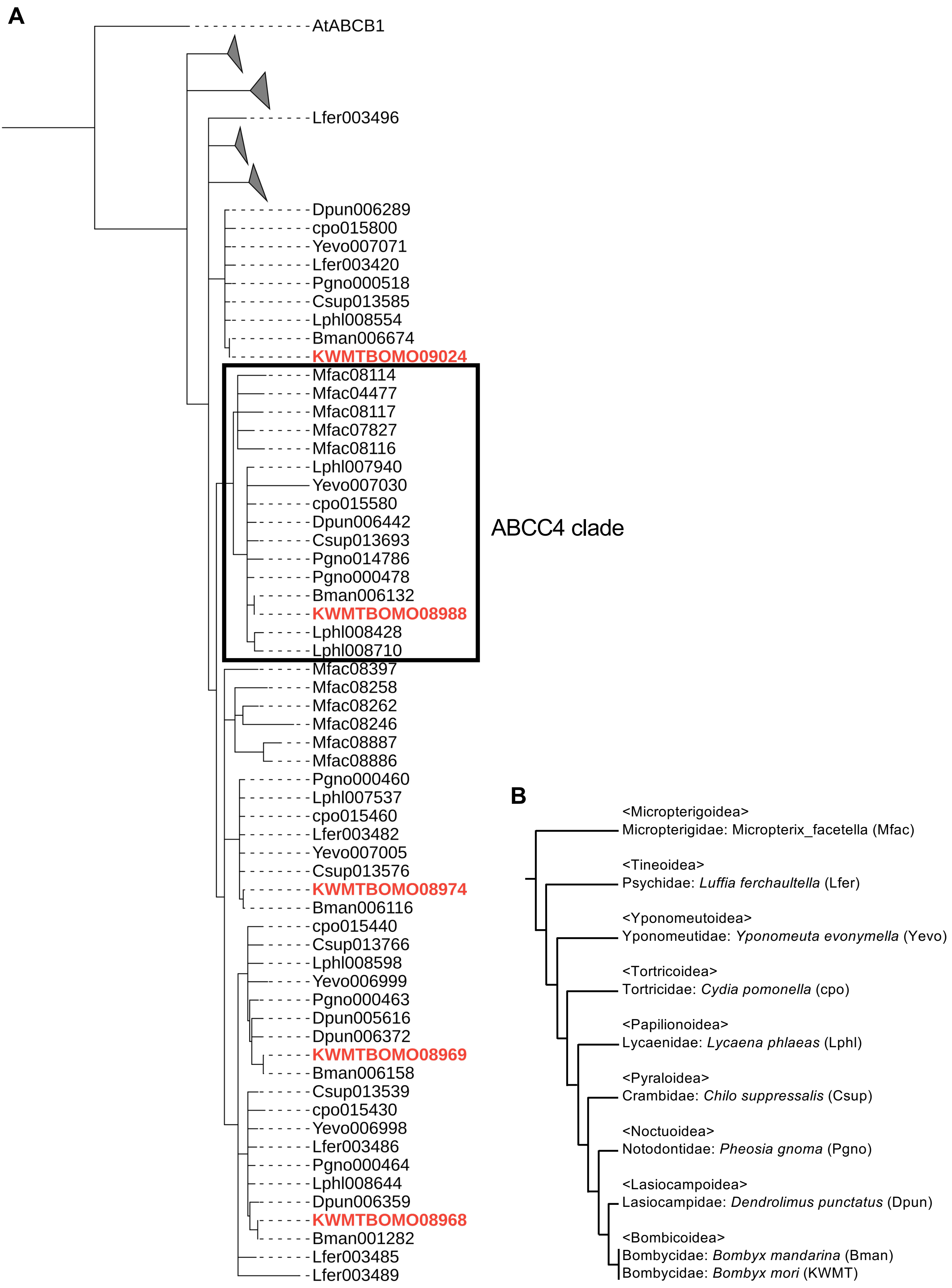


S1 Fig. Phylogenetic tree of lepidopteran-wide orthologous proteins of *Gc*-coded ABCCs
